# Cross-tissue Graph Attention Networks for Semi-supervised Gene Expression Prediction

**DOI:** 10.1101/2024.11.15.623881

**Authors:** Shiyu Wang, Mengyu He, Muran Qin, Yijuan Hu, Liang Zhao, Zhaohui Qin

## Abstract

High-throughput biotechnologies have significantly advanced precision medicine by enabling the exploitation of global gene expression patterns to enhance our understanding of disease etiology, progression, and treatment options. However, the tissue-specific nature of gene expression presents a challenge, particularly for less accessible tissues such as the brain, underscoring the need for computational methods to accurately impute gene expression in these critical but hard-to-reach tissues. While several attempts to impute gene expression in tissue-specific contexts have shown promising results, their reliance on regression analysis faces limitations due to the inability to capture complex, nonlinear relationships in gene expression patterns. In contrast, modern machine learning techniques, particularly graph neural networks, have demonstrated superior performance by efficiently modeling the intricate interactions among genes across different tissues. Therefore, we introduce gene expression imputation with Graph Attention Networks (gemGAT), a novel approach leveraging Graph Attention Networks (GATs) to enhance gene expression prediction across different tissues. gemGAT distinguishes itself by predicting the expression of all genes simultaneously, utilizing the full spectrum of genomic data to account for gene co-expressions and non-linear relationships. Validated through extensive experiments with Genotype-Tissue Expression (GTEx) data and a case study from the Alzheimer’s Disease Neuroimaging Initiative (ADNI), gemGAT demonstrates superior performance over existing methods by efficiently capturing non-linear gene co-expressions. This advancement underscores gemGAT’s potential to significantly contribute to precision medicine, showcasing its utility in advancing our understanding of gene expression in less accessible tissues.

## Introduction

Recent advances in high-throughput biotechnologies have made tremendous impact on biomedical sciences (Ashley 2016, Morash, Mitchell et al. 2018, Vilgis and Deigner 2018). As an example, global gene expression pattern has been exploited successfully to help scientists and clinicians to better understand the pathogenesis and progression of many human diseases (Golub, Slonim et al. 1999, Consortium 2015, Ahmed, Zeeshan et al. 2020). Despite the many successes, a fundamental challenge is that gene expression level is highly tissue-specific. It is likely that only gene expression pattern derived from disease-relevant tissues is informative for the understanding of human diseases (Maniatis, Goodbourn et al. 1987, Whitehead and Crawford 2005, Ong and Corces 2011). Except for few specimen such as whole blood, most disease-relevant tissues, such as the brain, are simply not accessible due to the invasiveness and cost. Alternative strategies must be taken. A promising strategy has emerged over recent years to overcome the challenge, which is to develop effective computational methods to accurately impute gene expression in inaccessible and disease-relevant tissues.

Over the years, multiple attempts have been made to tackle this important problem, yielding promising results. Despite the progress, significant limitations remain in the existing methods. Primarily, current methodologies predominantly rely on regression-based analysis(Halloran, Zhu et al. 2015, Xu, Liu et al. 2020, Basu, Wang et al. 2021) to perform the imputation. Such strategies are simple to implement and run fast. However, only capturing linear relationships is far from sufficient when modeling expression measures due to the diverse and sophisticated biological mechanisms that regulate the transcriptional programs in diverse tissues. Moreover, these methods primarily focus on modeling transcriptomics in a gene-by-gene basis, which is not only computationally intensive, but also inefficient to model complicated interactions among genes. On the other hand, modern machine learning-based approaches, such as graph neural networks (GNNs), is capable of modeling complicated nonlinear relationships among a large number of intricately correlated entities. These approaches have demonstrated superior performance (Hornik, Stinchcombe et al. 1989, Lu, Jin et al. 2021) and an enhanced ability to handle data interactions(Zhou, Cui et al. 2020, Wu, Cui et al. 2022).

Among various GNNs, Graph Attention Networks (GATs) offers a more nuanced and effective learning of node representations by learning the importance of each individual node based on their relationships and features(Velickovic, Cucurull et al. 2017). This capability aligns well with the task of cross-tissue gene expression imputation, where the impact of different genes on the prediction outcome varies. Therefore, in this work, we aim to leverage powerful GATs to overcome the limitations in existing methods in order to achieve accurate imputation of gene expression from observed source tissues (i.e., blood) to unobserved target tissues (i.e., brain). Here we introduce <u>g</u>ene <u>e</u>xpression i<u>m</u>putation with Graph Attention Networks (gemGAT), a novel machine learning approach that achieves accurate cross-tissue gene expression imputation by learning the intricate gene co-expression patterns present in both source and target tissues. Unlike existing methods, gemGAT has the capability to predict the expression of all genes simultaneously, offering a more efficient and comprehensive solution. Central to gemGAT is the use of GATs(Velickovic, Cucurull et al. 2017), which is able to capture both gene co-expressions and the potential non-linear, high-dimensional relationships among genes. Thereafter, by harnessing the power of deep neural networks, gemGAT exploits the full spectrum of genomic data, significantly advancing our ability to understand and predict gene expression in less accessible tissues.

To validate the efficacy of gemGAT, we conducted extensive experiments using gene expression data of multiple tissues from Genotype-Tissue Expression (GTEx) (Lonsdale, Thomas et al. 2013). We compare gemGAT with existing methods, highlighting its superior performance in predicting gene expression by emphasizing the importance of nonlinear gene co-expressions. To demonstrate the utility of gemGAT, we conduct a case study using Alzheimer’s Disease Neuroimaging Initiative (ADNI)(Mueller, Weiner et al. 2005) data, focusing on the practical application in disease gene discovery. Our experimental results not only underscore the advancements made by gemGAT but also highlight its potential to impact the field of precision medicine.

## Results

### Overview of gemGAT

The goal of gemGAT is to predict unobserved expression levels of genes in inaccessible tissue such as brain, referred to as the target tissue. To achieve this, two pieces of information are required as input: (1) gene expression data in an accessible tissue, referred to as the source tissue and (2) co-expression networks, referred to as networks, in both source and target tissues. See Figure 1, these are needed to enable gemGAT to leverage intricate correlation patterns among genes in modeling. Networks are assumed known *a priori* and are usually constructed by applying computational tools such as Weighted Gene Co-expression Network Analysis (WGCNA) (Langfelder and Horvath 2008) to extensive collections of multi-tissue gene expression data. Like gene expressions data, networks are also tissue-specific (Hu, Wan et al. 2010), meaning each tissue’s network is different from each other, often substantially. Even the number of genes in the network (defined to be genes connected to at least one other gene in the network), referred to as internal genes, differs dramatically across tissues. The proportion of internal genes in the genome ranges from 5% to 70%, with an average of 30%.

**Figure 1:**
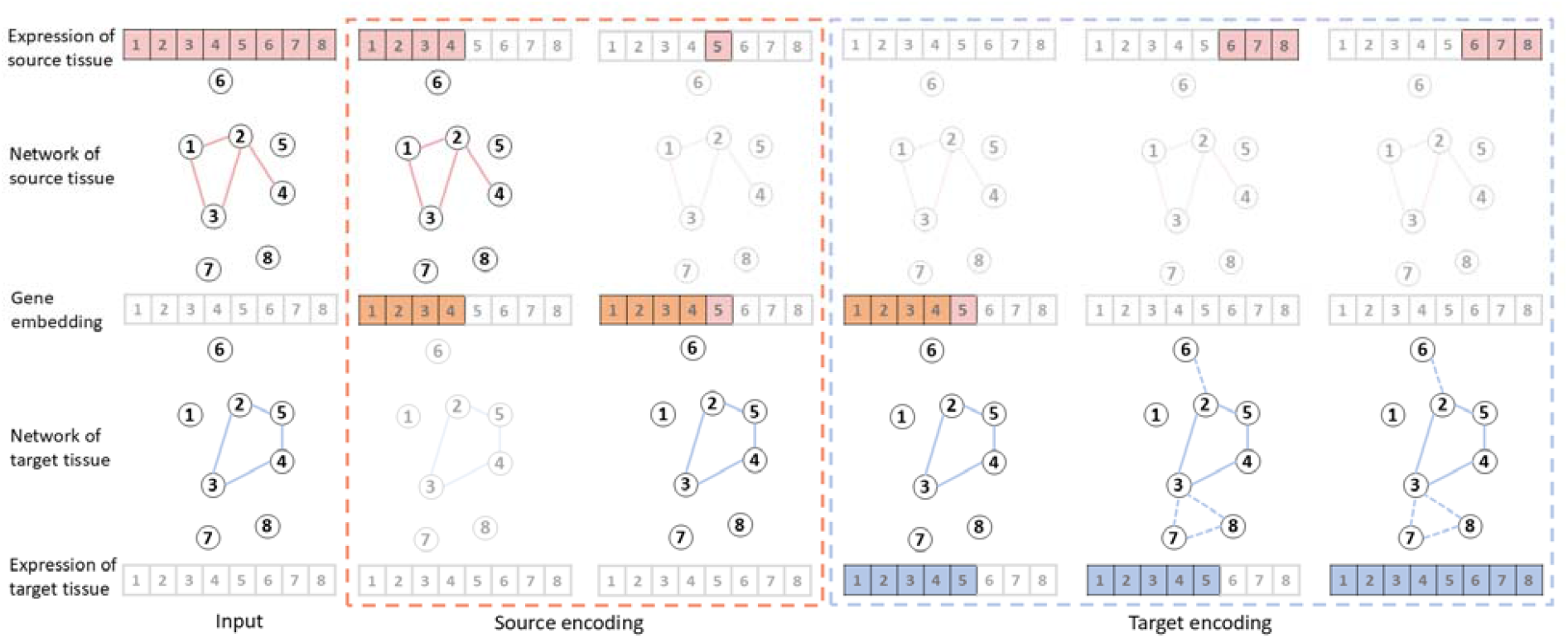
Illustration of the operation procedure of gemGAT. gemGAT predicts gene expressions in the target tissue (e.g., brain) using gene expressions observed in the source tissue (e.g., whole blood) and networks of both tissues. In the process of source encoding, embeddings of internal genes (i.e., gene 1-5) that are either in the network of the source tissue or in the network of the target tissue are computed. Those that are only internal genes of the target tissue (i.e., gene 5) are imputed by their expression in the source tissue. In the process of target encoding, embeddings of genes (i.e., gene 1-5) obtained in the previous step are used to predict their expressions in the target tissue. For those that are neither internal genes of the source nor the target tissue (i.e., gene 6-8), we first predict their connections with other genes (i.e., gene 1-5) using the existing network of the target tissue and the expressions of other genes in a semi-supervised manner. Then we predict their expressions leveraging the imputed gene-gene connections.

As illustrated in **Figure 1**, gemGAT operates in two stages: source encoding and target decoding.

#### Source encoding

The goal of this stage is to create an embedding of all the internal genes of either the source tissue or the target tissue. Both the gene expression data and the tissue networks are utilized. This is achieved by firstly utilizing GATs followed by a Multilayer Perceptron (MLP) to capture the structure of the network of source and target tissues to produce the embedding. Embedding of internal genes of the target tissue that are not internal genes of the source tissue is imputed by their expression levels in the source tissue.

#### Target decoding

In this stage, the expression levels of all the genes in the target tissue are predicted. This is achieved progressively in two steps. In the first step, we impute all the internal genes of either the source or the target tissue, accomplished by utilizing their embedding derived from the previous stage. This step uses GAT to capture the structure of the network of the target tissue, followed by a MLP for the purpose of prediction. In the second step, we impute the remaining genes, which are neither internal genes of the source nor the target tissue. This step is achieved by first imputing their connections with other genes using the existing network of the target tissue in a semi-supervised manner, and followed by predicting their expression levels leveraging the imputed gene-gene connections. Each process of the second step is formed by a similar GATs and MLP framework. The whole process is illustrated in **Figure 1**. More details of the gemGAT algorithm are provided in the **Methods** section. The implementation details have been illustrated in **Supplemental Materials A**.

### Performance evaluation on GTEx data

To demonstrate its performance, we test gemGAT on GTEx data, in which the whole blood is treated as the source tissue while 47 other tissues are treated as target tissues. GTEx data were split into mutually exclusive training and testing sets, where network construction and model training were conducted on the training set, and gene expression prediction was conducted on the testing set. More details can be found in the Methods section. Performance is measured using multiple metrics.

#### Gene-level correlation

In this part, we adopted the same metric used in Basu, Wang et al. (2021). Essentially, for each target tissue and for each gene, Pearson correlation coefficient (PCC) between the observed and the predicted across individuals is calculated. PCC helps us to assess whether the heterogeneity among individuals can be recapitulated in the predicted gene expression levels. **Figure 2A** shows the violin plots of gene-level PCC values for all 47 target tissues. We found that Brain Hippocampus, Brain Cerebellar Hemisphere, Adrenal Gland, Brain Substantia nigra and Brain Amygdala rank in the top according to the median PCCs. Five other brain tissues, Brain Hypothalamus, Brain Frontal Cortex (BA9), Brain Cortex, Brain Cerebellum and Brain Caudate (basal ganglia) rank at the bottom. Notably, we also observe that top ten and top 100 predicted genes according to their PCCs are quite different among brain tissues, as shown in **Figure S1** and **Figure S2** with an average Jaccard index of 0.13 and 0.18, respectively.

**Figure 2:**
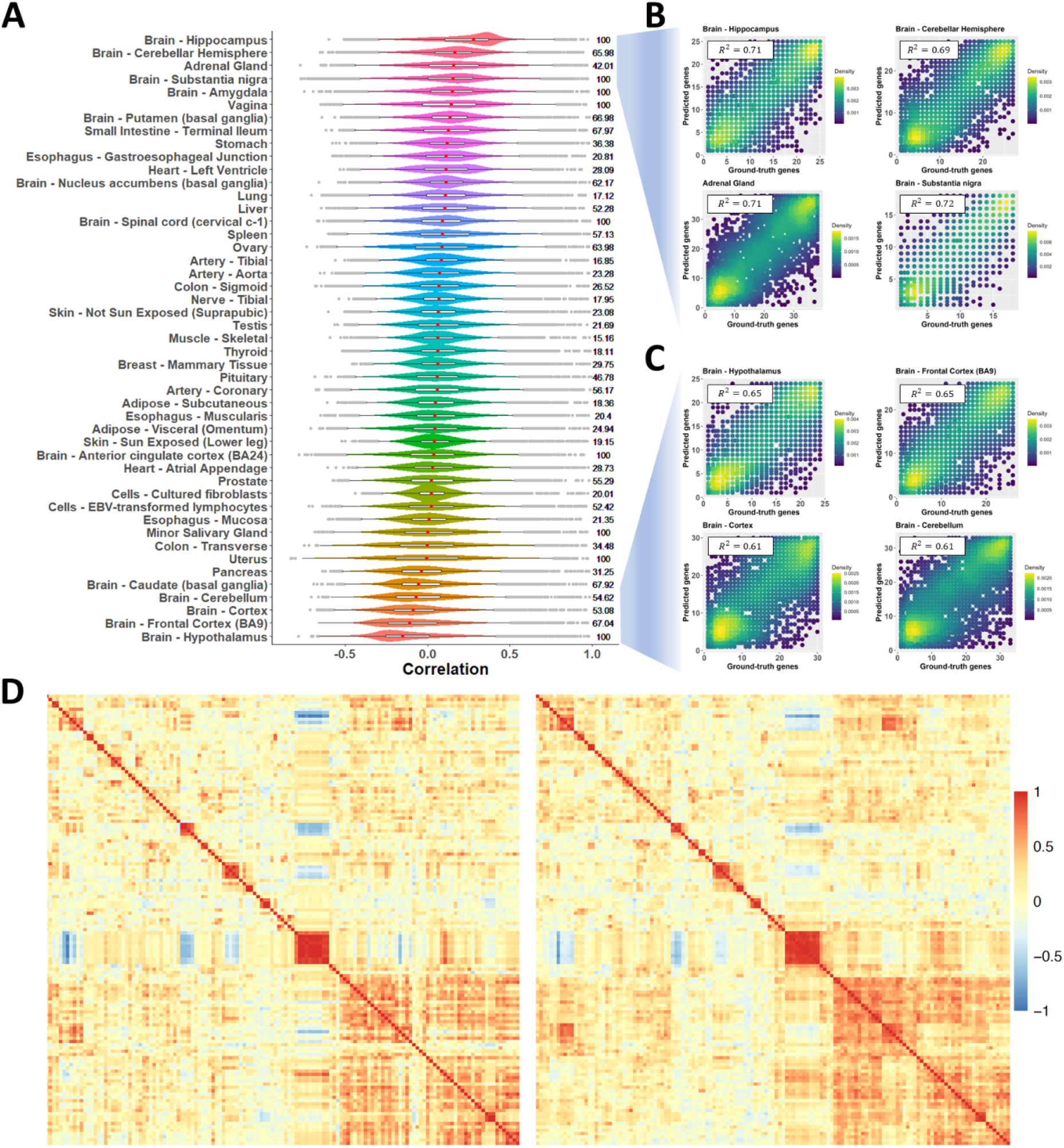
The predicting performance of gemGAT. **A**, Prediction accuracy in terms of Pearson correlation coefficient using gemGAT. Values on the right column indicate the percentage of genes whose predicted values are not significantly different from their true values, according to Wilcoxon signed rank test. **B**, Compare predicted genes against their true expressions in four best predicted tissues (i.e., Brain Hippocampus, Brain Cerebellar Hemisphere, Adrenal Gland and Brain Substantia Nigra). **C**, Compare predicted genes against their true expressions in four worst predicted tissues (i.e., Brain Hypothalamus, Brain Frontal Cortex (BA9), Brain Cortex and Brain Cerebellum). **D**, Heatmap that shows correlation among accurately predicted genes (left) in Brain Hippocampus and the correlation among their true values in the same tissue (right).

**Figure 3:**
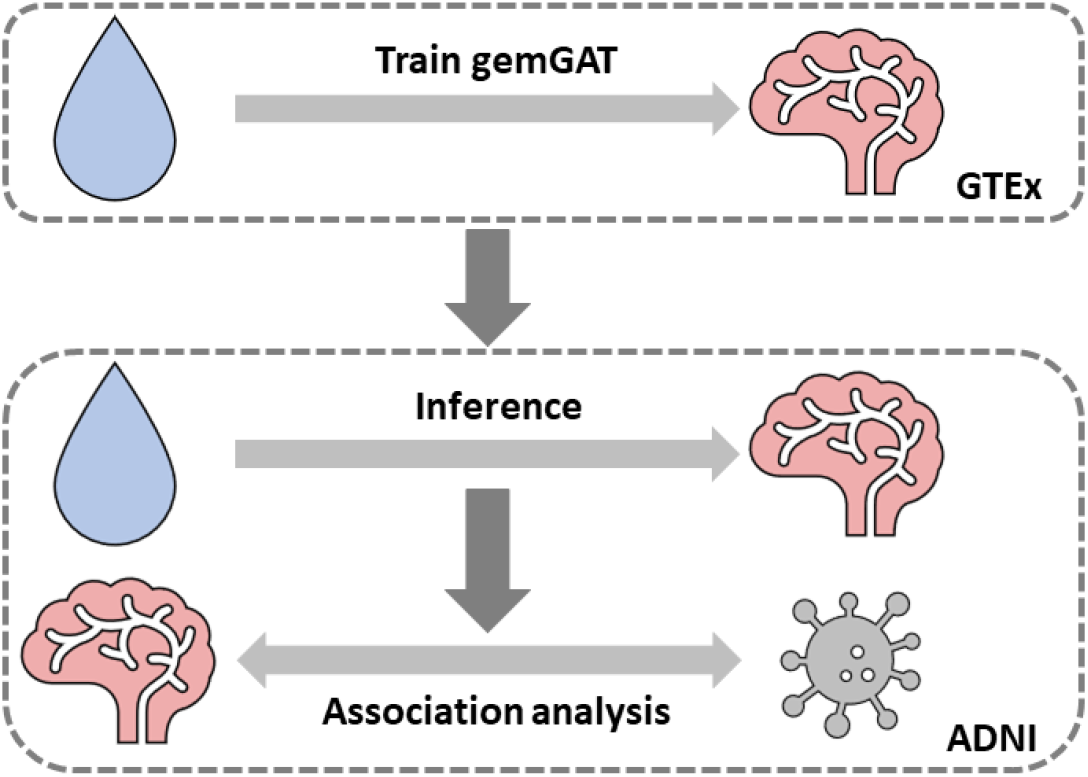
Design of case study. gemGAT is trained in GTEx dataset first to predict gene expressions in the target tissue (e.g., brain) from those in the source tissue (e.g., whole blood). Then the trained model is applied to the source tissue in ADNI dataset to predict gene expressions in the same target tissue but for individuals in ADNI dataset. Next, the predicted gene expressions are employed for association analysis to identify AD-related genes.

In addition to global correlation patterns, we also zoom in to selected genes for a close up. Specifically, we present scatter plots of rank orders among donors for observed and predicted gene expression levels of selected genes. Top 100 genes in the top four tissues (i.e., Brain Hippocampus, Brain Cerebellar Hemisphere, Adrenal Gland and Brain Substantia nigra) are plotted in **Figure 2B** and the bottom four tissues (i.e., Brain Hypothalamus, Brain Frontal Cortex (BA9), Brain Cortex and Brain Cerebellum) are presented in **Figure 2C**. Among these top genes, *R*^2^ values are 0.71, 0.69, 0.71 and 0.72, respectively for these four tissues, indicating strong correlation between predicted gene expression levels and the observed. For the bottom four tissues, we can still observe a robust correlation between the observed and the predicted with *R*^2^ of 0.65, 0.65, 0.61 and 0.61, respectively. **Figure 2A, B and C** together indicate that gemGAT is able to consistently achieve high-quality predictions for all tissues. The summary of all the per-gene correlation measures in all tissues can be found in **Supplemental Materials C**.

At the single gene level, we select a few genes as representatives and plot the predicted expression levels again the observed expression levels in a scatter plot for each of the gene (Supplementary **Figures S3** and **S4**). We used the aforementioned four top tissues and four bottom tissues in this part. Five genes were selected for each tissue, with varying levels of observed-predicted correlations (1 percentile, 20 percentile, 40 percentile, 60 percentile, 80 percentile). Our results demonstrated that predicted values and true expressions show strong correlation at the individual gene level for top genes (1 percentile and 20 percentile), regardless of top tissues or bottom tissues.

#### Correlation matrix

For accurate prediction, it is important to ensure that intricate relationships among genes are preserved as much as possible. Gene-gene relationships can be represented by their correlation matrix. From **Figure 1**, we know that it is unrealistic to require accurate prediction for every gene in the genome. Therefore, we focus on genes that are “well-predicted”, defined as those with *PCC* > 0.7 between observed and predicted. Using the Brain Hippocampus tissue as an example, there are 536 genes meet this threshold. We plot the correlation matrix among these genes using both predicted values (**Figure 2D left**) and observed values (**Figure 2D right**). We found that the two heatmaps resemble each other nicely, indicating that gemGAT is able to recover the correlation relationship among different genes.

### Application to identify disease-associated genes

We conduct a case study to evaluate the effectiveness of predicted expression values in inaccessible but disease-relevant tissues. Specifically, we aim to identify differentially expressed genes (DEGs) and pathways in Brain Hippocampus, in a study of Alzheimer’s disease (AD) under the case-control study setting. Brain Hippocampus is among the earliest affected brain regions for AD patients and its dysfunction is believed to underlie the core feature of AD-memory impairment (Maruszak and Thuret 2014). In the clinic, it is almost impossible to have access to brain tissues from live patients.

Therefore, it would be highly desirable to predict gene expression in disease-relevant brain regions such as hippocampus based on expression values measured in the most clinically accessible tissue, e.g., whole blood. To test this idea, we apply gemGAT to gene expression data from Alzheimer’s Disease Neuroimaging Initiative (ADNI), which is collected from the whole blood of 188 AD patients and 634 normal individuals (Note that we merge “Normal” and “Mild Cognitive Impairment” individuals together as the “Normal” cohort).

#### DEG analysis

After applying gemGAT, we run logistic regression on the predicted expression levels to identify DEGs between AD patients and normal controls(Choi, Labadorf et al. 2017). We used p-value of 0.05 as the significance threshold and permutation test (Camargo, Azuaje et al. 2008) is performed on those significant genes to account for multiple comparisons.

**Table 1** lists the top genes showing the most significant changes in predicted gene expression values between AD patients and controls. Remarkably, we found that many of the top-ranked genes are indeed AD-associated. For instance, the top gene on the list--*CLDN20* (p-value= 1.34 ×10^−4^), is a member of the claudin family that has been previously found to be differentially expressed in AD brains (Günzel and Yu 2013). In another paper, Spulber, Bogdanovic et al. (2012) showed that claudin expression profile separates AD cases from vascular dementia cases. For the second gene on the list— *COBL* (p-value=1.56×10^−4^), in a recent genome-wide association study (GWAS) conducted in African Americans on AD by Mez, Chung et al. (2017), a variant located upstream of this gene achieves genome-wide significant association. The third gene on the list—*CSNK1D*, (p-value=2.53×10^−4^), has been found to be upregulated in AD brain (Yasojima, Kuret et al. 2000), which is consistent with our predicted profile (regression effect size=3.66). The fourth gene on the list is *EGR1* (p-value=7.06×10^−4^). In a recent study, this gene was found to play a key role in keeping the brain cholinergic function intact in the preclinical stages of AD(Hu, Chen et al. 2019). The fifth gene on the list is *FBXL13* (p-value=8.64×10^−4^), Floudas, Um et al. (2014) found evidence of involvement of *FBXL13* in late-onset AD. *FBXL13* encodes a protein belonging to the F-box protein family. Members of this family take part in SKP1-CUL1-F-box (SCF) protein complexes that act as protein-ubiquitin ligases(Maglott, Ostell et al. 2010). The ubiquitin-proteasome system is involved in protein turnover and degradation and is perturbed in AD(Floudas, Um et al. 2014). More discussion on those AD-associated genes is in **Supplemental Materials D**. The significance of the association between these genes and AD are also reiterated by the fact that all of their permutation p-values are lower than 0.05.

**Table 1:** Top ten AD-associated genes in Brain Hippocampus predicted by gemGAT based on logistic-regression p-values.

| Rank | Gene | p-value | perm. p-value | References |
| --- | --- | --- | --- | --- |
| 1 | *CLDN20* | $1.34 \times 10^{-4}$ | $2.00 \times 10^{-4}$ | (Romanitan, Popescu et al. 2010, Spulber, Bogdanovic et al. 2012) |
| 2 | *COBL* | $1.56 \times 10^{-4}$ | $1.00 \times 10^{-4}$ | (Mez, Chung et al. 2017) |
| 3 | *CSNK1D* | $2.53 \times 10^{-4}$ | $2.00 \times 10^{-4}$ | (Schwab, DeMaggio et al. 2000, Yasojima, Kuret et al. 2000) |
| 4 | *EGR1* | $7.06 \times 10^{-4}$ | $1.00 \times 10^{-3}$ | (Zhu, Unmehopa et al. 2016, Hu, Chen et al. 2019) |
| 5 | *FBXL13* | $8.64 \times 10^{-4}$ | $2.00 \times 10^{-4}$ | (Watanabe, Von Der Kammer et al. 2012, Floudas, Um et al. 2014) |
| 6 | *HHLA2* | $1.01 \times 10^{-3}$ | $1.60 \times 10^{-3}$ | (Sabaie, Tamimi et al. 2023) |
| 7 | *IGSF6* | $1.27 \times 10^{-3}$ | $4.00 \times 10^{-4}$ | (Hargis and Blalock 2017) |
| 8 | *SCIMP* | $1.32 \times 10^{-3}$ | $2.00 \times 10^{-3}$ | (Jansen, Savage et al. 2019, Andrade-Guerrero, Santiago-Balmaseda et al. 2023) |
| 9 | *VEGFC* | $1.36 \times 10^{-3}$ | $1.20 \times 10^{-3}$ | (Kalaria, Cohen et al. 1998, Hohman, Bell et al. 2015) |
| 10 | *WLS* | $1.67 \times 10^{-3}$ | $1.40 \times 10^{-3}$ | (Aghaizu, Jin et al. 2020, Martínez and Inestrosa 2021) |

For comparison, we also conducted DEG analysis directly on observed expression of the same set of genes from whole blood. The top-ranked DE genes in whole blood are summarized in **Table S1**. Among the top ten genes, three genes—*FBXL13, WLS* and *IGSF6* also appeared in the top ten genes identified in Brain Hippocampus. For the remaining genes, searching the existing literature, we found only three of them are AD-related.

Between the two sets of results, as expected, DEGs identified in blood (observed) are largely distinct from DEGs identified in the hippocampus (predicted). To be specific, as Wilcoxon signed-rank test conducted to compare the two sets of genes ranked by their association with AD returned a p-value of 4.11×10^−11^

#### Pathway analysis

In addition to DEG analysis, we also conduct gene set enrichment analysis (GSEA)(Subramanian, Tamayo et al. 2005) on the 186 KEGG pathway (Kanehisa 2002) using “fgsea()” in R 4.2.0. Interestingly, we observe multiple AD-related pathways show up in our result, as shown in **Figure 4 A**.

**Figure 4:**
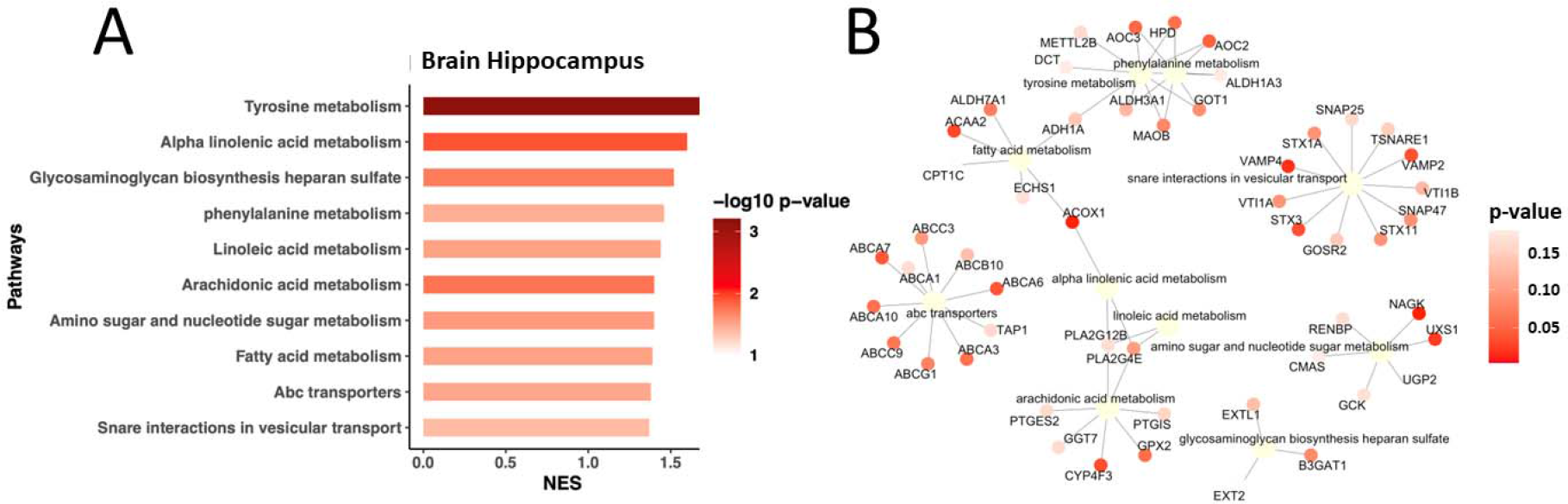
Pathways and identified associated with AD based on predicted genes. **A**, the top-ranked ten AD-associated pathways by NES (i.e., normalized enrichment score) from GSEA based on predicted genes in Brain Hippocampus. **B**, Visualization of pathways identified in A by showing genes with DE p-values smaller than 0.2.

Among the two sets of top ten most enriched pathways identified in Brain Hippocampus and whole blood, respectively, as shown in **Figure 4 A, Table 2** and **Table S2**, only one pathway (fatty acid metabolism) is found in both sets. All top ten enriched pathways identified in Brain Hippocampus are supported by the existing literature (**Table 2**). For instance, the expression of +tyrosine metabolism (p-value= 7.02×10^−4^) is found to be significantly different between AD patients and controls(Forssell, Eklöf et al. 1989). Alpha linolenic acid metabolism (p-value=1.23×10^−2^) regulates amyloid precursor protein processing by mitogen-activated protein kinase pathway and neuronal apoptosis in amyloid beta-induced SH-SY5Y neuronal cells, and thus may be a beneficial agent for promoting prevention of AD(Lee, Lee et al. 2018). Glycosaminoglycan biosynthesis heparan sulfate (p-value= 1.88×10^−2^) is supported by a study that found that the nuclei of selected neurons and a small number of microglia are immunopositive for heparan sulfate glycosaminoglycan in contrast to controls(Su, Cummings et al. 1992). Phenylalanine metabolism (p-value= 3.45×10^−2^) is dysregulated in human hippocampus with AD-related pathological changes(Liu, Yang et al. 2021). Besides, an excessive intake of linoleic acid (p-value= 2.99×10^−2^) are found to lead to the formation of oxidized linoleic acid metabolites, which have been associated with AD(Mercola and D’Adamo 2023). Furthermore, when plotting those top-ranked pathways (**Figure 4B**), we observe that several key AD-related genes are included. For instance, ACOX1, shared by fatty acid metabolism and alpha linolenic acid metabolism, is found to be related to microglial peroxisomal dysfunction, which directly and indirectly affects the inflammatory response of microglia in the brain(Uzor, McCullough et al. 2020). Furthermore, the accumulation of amyloid *β* peptide (A*β*) in the brain of AD patients begins many years before clinical onset(Belfiori-Carrasco, Marcora et al. 2017). The gene HPD that is shared by tyrosine metabolism and phenylalanine metabolism is identified as the suppressor of intraneuronal accumulation of A*β* (Belfiori-Carrasco, Marcora et al. 2017). Another two genes, GOT1 and MAOB, that are shared by the same pathways are also found to play key roles in AD pathogenesis. GOT1 is found to be downregulated in neurodegenerative diseases such as AD(Saxena, Murthy et al. 2021), and MAOB is elevated in AD neurons and is able to regulate A*β* level(Schedin-Weiss, Inoue et al. 2017). Also, a significant decrease in PLA2G4E is evident in the brain of late-stage AD patients(Pérez-González, Mendioroz et al. 2020), which is shared by three top-ranked pathways (i.e., linoleic acid metabolism, alpha linolenic acid metabolism and arachidonic acid metabolism) based on **Figure 4B**.

**Table 2:**
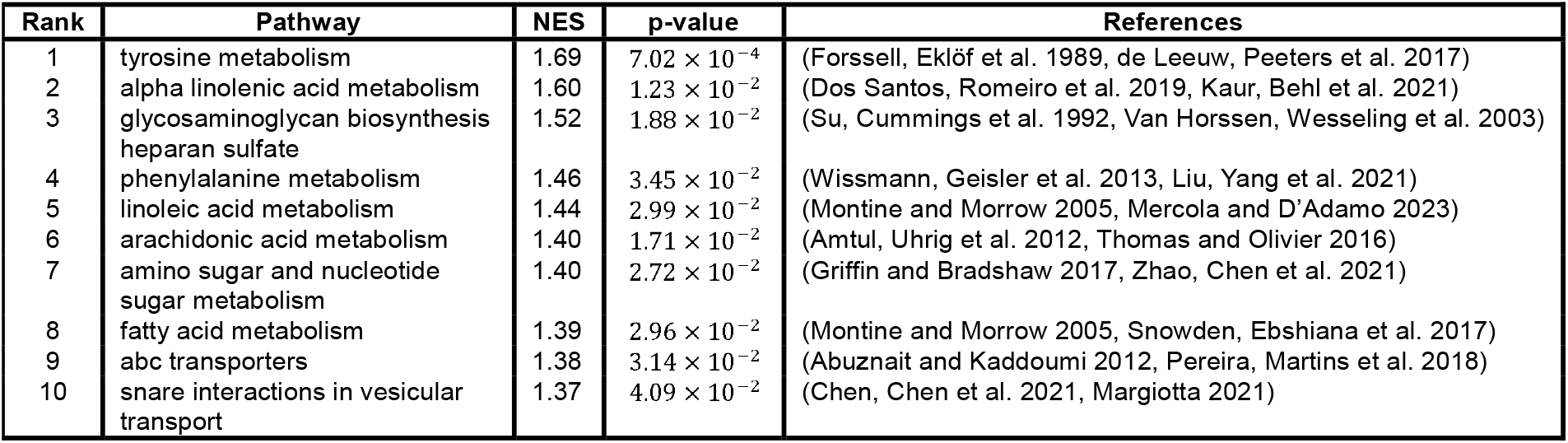
Top ten pathways identified by applying GSEA to p-values of differential expression analysis on predicted genes by gemGAT.

In contrast, not all of the top ten pathways found in whole blood are AD-related. Moreover, the normalized enrichment score (NES) of pathways enriched in the Brain Hippocampus (1.47 on average) are higher that the NES scores of pathways in the Whole Blood (1.34 on average), with p-value 0.02 (Wilcoxon Signed Rank Test). Also, enrichment of all top ten pathways in Brain Hippocampus illustrated in Table 2 are statistically significant based on their permutation p-values, whereas only two out of top ten enriched pathways in Whole Blood are significant. Notably, the shared pathway fatty acid metabolism has the permutation p-value of 7.95×10^−2^ Our results are consistent with the fact that AD is a neurodegenerative disorder that shows disruption in selected brain regions such as hippocampus (Tanaka, Shiojiri et al. 1989, McGeer, Kawamata et al. 1992, Mirza, King et al. 2017).

In conclusion, when applying gemGAT to predict gene expression levels in AD-relevant brain tissues, subsequent DEG analysis reveals many of the most enriched pathways are AD-related which is supported by the existing literature, underscoring the effectiveness of the proposed GAT-based approach. This could also boost the usage of gemGAT and provide the community more insights into biological and pathological processes of those biological evidence.

### Comparison with regression-based methods

To evaluate the performance of gemGAT, we compare it with two competing methods for cross-tissue gene expression prediction.

TEEBoT is a linear model-based approach, which incorporates the principal components of whole blood gene expression along with sample covariates (e.g., sex, age, etc), and principal components of eQTL single-nucleotide polymorphisms (eSNP) and whole blood splicing profile (WBSp) to predict gene expressions of other tissues(Basu, Wang et al. 2021). Predicting each gene requires training one TEEBOT.

PrediXcan is a gene-based association method aims to predict the component of gene expression influenced by an individual’s genetic profile and correlates the “imputed” gene expression profile with the phenotype(Gamazon, Wheeler et al. 2015). We apply an embedded model of PrediXcan for the imputation of gene expressions from SNPs. Specifically, this approach fits one linear model in each tissue to predict gene expression of a specific gene solely based on SNP genotype profile. Similar to TEEBOT, this process requires training a unique model for each individual gene, and model involves the number of reference alleles of all SNPs, whose dimension is typically much higher than the principal components employed to fit TEEBOT.

As shown in Figure **5A**, when compare gemGAT with competing methods using median Kendall rank correlation coefficient (KCC), we found gemGAT predominantly outperforms the two competing methods in 39 out of 47 tissues (i.e., 83%), while TEEBOT performs the best in the remaining eight tissues. We also evaluate gemGAT against competing methods based on the number of genes for which they can make accurate predictions (i.e., *PCC* > 0.7). As shown in **Figure 5B**, we find that gemGAT makes more accurate predictions in 43 out of 47 tissues (i.e., 91.49%). The training process of competing methods involves dealing with high-dimensional eSNPs or WBSp, which becomes computationally intensive when there are thousands of genes to be predicted. Additionally, non-linear relationships among genes cannot be fully captured by simpler linear models.

**Figure 5:**
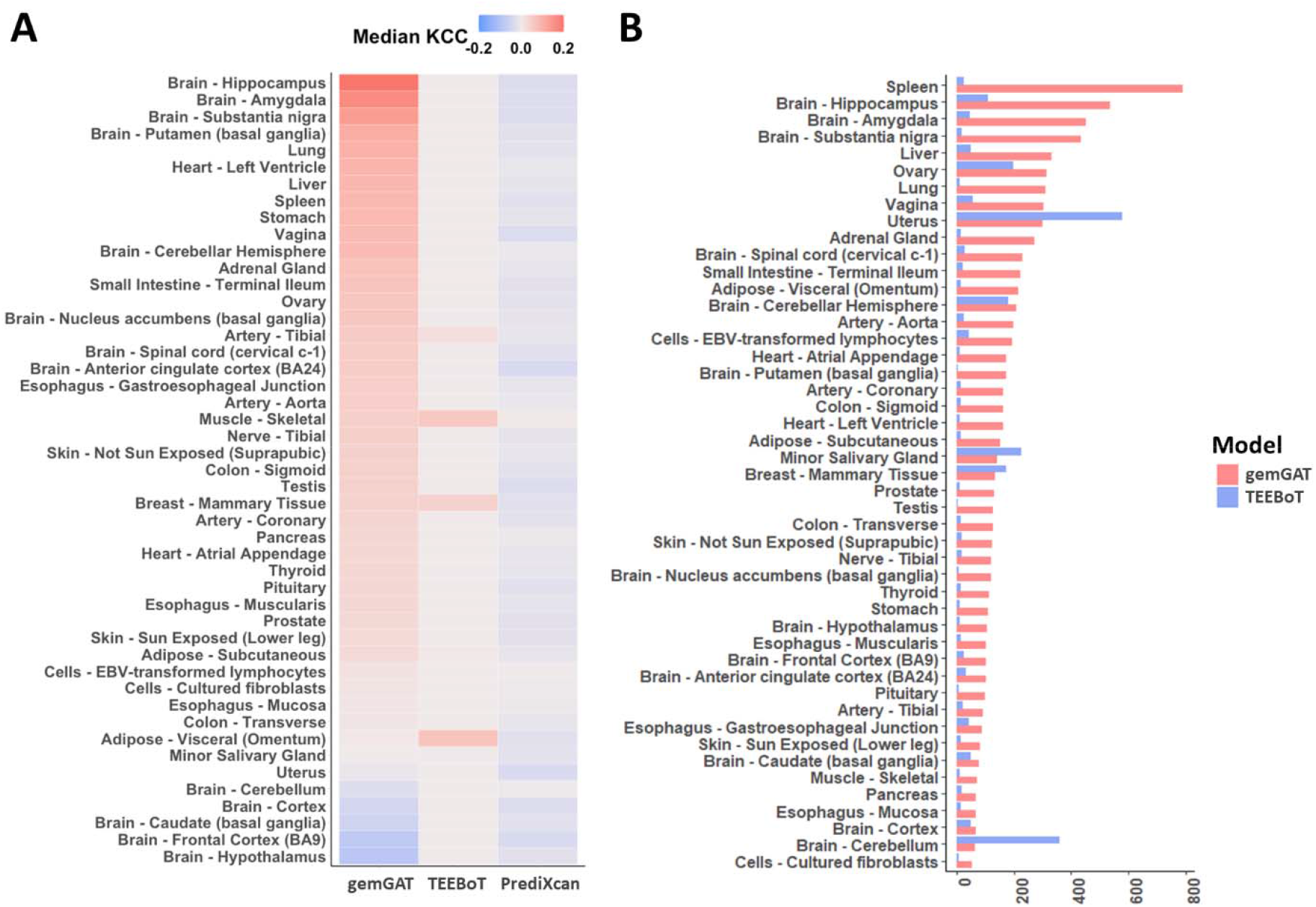
Comparing predicting capability with existing methods. **A**, Predicting accuracy in terms of KCC using gemGAT, TEEBOT and PrediXcan. **B**, Number of well-predicted genes in terms of PCC (i.e., with PCC>0.7) using gemGAT and TEEBOT.

In addition to compare predicted versus observed correlation, in the real-world applications, such as differential expression analysis, the overall distribution of the expression levels of a gene draws more attention is arguably more important than their actual values. Therefore, we also assess the proposed model by investigating if the overall distribution of genes could be preserved by the model. **Figure S5** shows the proportion of predicted genes in each tissue that preserve their original distributions according to Kolmogorov–Smirnov (KS) test (i.e., p-value>0.5). The results suggest that gemGAT outperforms TEEBOT in 43 out of 47 tissues (i.e., 91%). Across all sample for each gene, versus across all genes for each sample.

## Methods

In this section, we explain detailed techniques that leveraged by gemGAT.

### Dataset and gene expression data processing

We obtain tissue-specific gene expression data from the Genotype-Tissue Expression (GTEx) project(Lonsdale, Thomas et al. 2013) using the GTEx portal^1^. Gene expression data from 47 tissues (excluding two tissues with insufficient sample size) were included in our study. To improve the predictive power, we exclude lowly-expressed genes, which are defined as those with less than two read counts across all subjects in a specific tissue(Wang, Liu et al. 2021). We then pair gene expressions in whole blood with the other 46 tissues one-by-one as source-target pairs, removing subjects and genes if there is missing data in either tissue. Subjects in each whole blood-target tissue pair are split into training and testing sets in the ratio of 80% and 20%, respectively.

### Gene co-expression network construction

After splitting all the expression data into training and testing, we conducted Weighted Correlation Network Analysis (WGCNA) on the training data using the “WGCNA” package in R 4.2.0(Langfelder and Horvath 2008) on all 46 tissues. For each tissue, we used the “pickSoftThreshold()” function to calculate the scale-free topology fitting index for different soft thresholding powers and selected the appropriate power value to construct the correlation networks. Then, we used the “adjacency()” function to compute the adjacency matrix by transforming the Pearson correlation coefficients between each pair of genes into adjacency values using the chosen power value. Finally, we used a predefined threshold *δ* to determine the edges among genes in the network. In such networks, genes are represented as nodes and the edges represent the presence of correlations between the two genes(Langfelder and Horvath 2008). Note that in practice, such networks are assumed to be fixed and known *a priori*.

### Overview of gemGAT

Let 𝒢_*s*_ and 𝒢_*T*_ be gene-gene networks of the source and the target tissue, respectively, and 𝒱_*s*_ and 𝒱_*T*_ be sets of genes contained in 𝒢_*s*_ and 𝒢_*T*_. Usually, 𝒱_*s*_ ≠ 𝒱_*T*_ and 𝒱_*s*_, 𝒱_*T*_ ⊂ 𝒱, where 𝒱 is the collection of all genes in the genome. As shown in **Figure 1**, we first conduct source encoding to encode genes in the source tissue leveraging 𝒢_*s*_. Gene embeddings generated in this step carry the information of genes in the source tissue. In the second step, during target decoding, we firstly focus on predicting genes that are in networks of either source or target tissues: 𝒱_*in*_ =𝒱_*s*_ ∪ 𝒱_*T*_. Specifically, we employ Graph Attention Networks (GATs)(Velickovic, Cucurull et al. 2017) to incorporate topological information found in both networks followed by Multilayer perceptron (MLP) as the prediction layer. Then, for the remaining gene not in 𝒱_*in*_ (or genes in 𝒱\ 𝒱_*in*_), we intend to impute their interactions with other genes and among themselves via semi-prediction. Lastly, we leverage the imputed network to predict expressions of genes not in 𝒱_*in*_ (or genes in 𝒱\𝒱_*in*_). supervised link prediction. Lastly, we leverage the imputed network to predict expressions of genes not in 𝒱_*in*_ (or genes in 𝒱\𝒱_*in*_).

### Source encoding

First of all, we encode genes in 𝒱_*s*_ while leveraging gene-gene networks 𝒢_*s*_ in the source tissue using GATs:

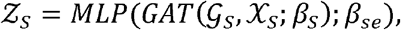

where χ_*s*_ is the expression of genes in 𝒱_*s*_ and *β*_*s*_ is the model parameter. The detailed formulation of *GAT*(·)is in **Supplemental Materials G**. We further merge 𝒢_*T*_ with other genes in 𝒱_*in*_ to 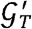 as the gene-gene network that contains all genes in 𝒱_*in*_, in which those newly-added genes only connect to themselves. As 𝒵_*s*_ is the embedding of genes only in 𝒱_*s*_, to leverage 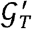,we impute the embedding of genes that in 𝒱_*in*_ but not in 𝒱_*s*_ those newly-added genes only connect to with their expression in the source tissue.

### Target decoding

#### Predict genes in 𝒱_in_

Finally, for all genes in 𝒱_*in*_, we predict their expression in the target tissue as follows:

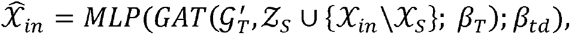

where χ_*in*_ is the expression of genes in 𝒱_*in*_ in the source tissue, and *β*_*T*_ and *β*_*td*_ are two sets of model parameters of GATs and a MLP, respectively. The detailed formulation of *MLP*(·) is in **Supplemental Materials H**. Until now, we have predicted expression of all genes in 𝒱_*in*_ within gene-gene networks of source and target tissues. Nevertheless, there are still genes in 𝒱 but not in 𝒱_*in*_, the expression of which has not been predicted in the target tissue. Those genes are not in 𝒱_*s*_ or 𝒱_*T*_, so that directly leveraging gene-gene networks, 𝒢_*s*_ or 𝒢_*T*_ to predict their expression becomes challenging.

#### Semi-supervised link prediction

As the remaining genes to be predicted are not in 𝒱_*s*_ or 𝒱_*T*_, we intend to impute their interactions both with other genes and among themselves to leverage gene-gene networks in the prediction task. As shown in **Figure 1**, this is naturally formed into a graph link prediction task(Berton, Valverde-Rebaza et al. 2015, Hisano 2018). Specifically, we merge 𝒢_*T*_ with other genes in 𝒱 to initialize 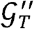 as the network that involves all genes in 𝒱. Newly-added genes 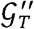 to compared to 𝒢_*T*_ only connect to themselves at the beginning, whereas we aim to predict their interactions with other genes. We keep leveraging GATs and borrow predicted expressions 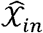 to serve as the input of genes in 𝒱_*in*_. For genes in 𝒱\𝒱_*in*_, we use their expressions in the source tissue χ \χ_*in*_ as the input, where χis the set of gene expressions of all genes in 𝒱. Specifically, the imputed network in the target tissue is obtained by the following process.

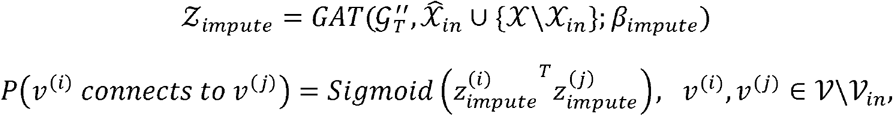

where 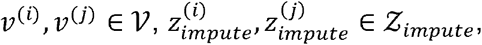 and *β*_*impute*_ is the model parameter of the GATs. That is to say, we assume that the interaction between two genes follows the Bernoulli distribution with the probability computed by the dot product of their embeddings. During the training process, we keep updating 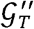 following the above process by adding newly imputed interactions among genes. In the end, powered by or GATs, the gene-gene network in the target tissue grows as the complementary of the original 𝒢_*T*_, the construction of which might be limited to the prior knowledge or traditional methods (e.g., WGCNA). 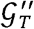 provides topological and biological insights for the prediction of genes in 𝒱\𝒱_*in*_.

#### Predict genes in 𝒱\𝒱_in_

Once the gene-gene network 𝒢_*T*_ in the target tissue is imputed to 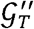 following the method above, as shown in **Figure 1**, we predict expressions of genes in 𝒱\𝒱_*in*_ by leveraging their interactions in 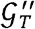.Specifically, the out-network prediction is conducted in the following manner.

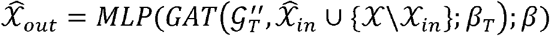

The predicting results 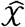 for all genes in 𝒱 is obtained:

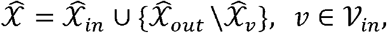

where 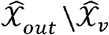 is the output results in 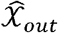 for genes in 𝒱_*in*_.

#### Learning objective and training details

During training, we employ the Mean Squared Error (MSE) between predicted expressions and true expressions to serve as reconstruction loss:

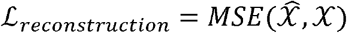

Furthermore, we borrow Binary Cross Entropy (BCE) as the reconstruction loss of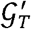:

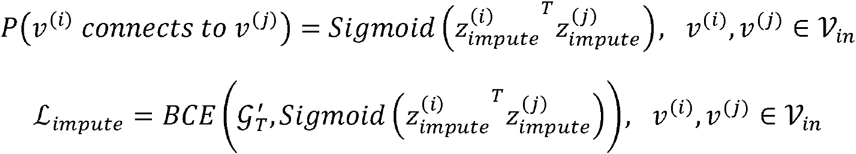

The final learning objective is composed of both ℒ_*reconstruction*_ and ℒ_*impute*_:

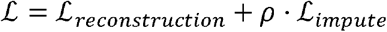

The training process is performed on the 64-bit machine with an NVIDIA GPU, NVIDIA GeForce RTX 3090. The total training process takes up to 300 epochs until converge using PyTorch 1.8.1 on Python 3.9.

#### Competing methods

To demonstrate the effectiveness of the proposed model, we compare it with two competing methods.

In a recent study, Basu, Wang et al. (2021). developed a linear regression-based method named TEEBoT to predict gene expression levels across tissues. To reduce dimension, TEEBoT utilizes estimated principal components (PCs) instead of actual expression values in the regression models. The potential drawback of TEEBoT is that it can only capture linear correlated between genes, while the potential non-linear relationship among gene expressions caused by complex biological process is under-explored.

PrediXcan is a computational method aims to estimate the component of gene expression determined by an individual’s genetic profile (Gamazon, Wheeler et al. 2015). Although it is not designed for imputing tissue-specific gene expression levels, it is nevertheless capable of imputing tissue-dependent gene expression levels using the linear model trained with genome-wide variation profiles.

### Evaluation metrics

We use Pearson correlation coefficient (PCC) calculated between predicted and actual gene expression levels to evaluate the performance of the prediction algorithm. Let *g*_1_ and *g*_2_ be two vectors of *N* matched gene expressions (e.g., 𝒢_1_ and 𝒢_2_ are predicted and original gene expressions, respectively). Then,

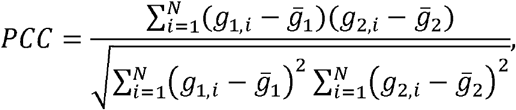

where *g*_1,*i*_ is the i-th gene in *g*_1_ and *g*_2,*i*_ is the i-th gene in 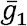 and 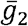 are mean values of *g*_1_ and *g*_2_, respectively.

When compare gemGAT with baselines, to deal with tied values, we switch from PCC to Kendall rank correlation coefficient (KCC) calculated between predicted and actual gene expression levels to evaluate the performance of the prediction algorithm. Let 𝒢_1_ and 𝒢_2_ be two sets of *N* matched gene expressions (e.g., 𝒢_1_ and 𝒢_2_ are predicted and original gene expressions, respectively). Then,

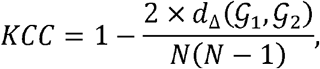

Where *d*_Δ_(𝒢_1,_𝒢_2_) is the number of different pairs between 𝒢_1_ and 𝒢_2_.

### Verification of AD-associated genes by permutation test

We perform permutation test to empirically validate the significance of identified DEGs. Specifically, we conduct logistic regression of AD against predicted gene expressions to identify DEGs. For a particular DEG *g*, we record its rank *r*_*g*_ among all genes based on their p-values. Then, we randomly permute disease labels for 1,000 times, and each time we conduct logistic regression that returns p-values of AD association and we record the rank of the same gene *g*:{*r*_1,_ *r*_2,…_ *r*_1000_}. Finally, we compute the permutation p-value as follows:

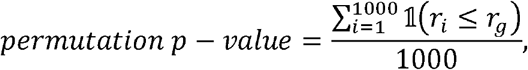

Where 𝟙(*r*_*i*_ ≤ *r*_*g*_) = 1 if *r*_*i*_ ≤ *r*_*g*_, otherwise 𝟙(*r*_*i*_ ≤ *r*_*g*_) = 0.

## Software

The implementation of gemGAT is available in GitHub repository: https://github.com/shi-yu-wang/gemGAT.

## Footnotes

1 https://gtexportal.org/home/

## Reference

Abuznait, A. H. and A. Kaddoumi (2012). “Role of ABC transporters in the pathogenesis of Alzheimer’s disease.” ACS chemical neuroscience 3(11): 820–831.

Aghaizu, N. D., et al. (2020). “Dysregulated Wnt signalling in the Alzheimer’s brain.” Brain Sciences 10(12): 902.

Ahmed, Z., et al. (2020). “Human gene and disease associations for clinical-genomics and precision medicine research.” Clinical and translational medicine 10(1): 297–318.

Amtul, Z., et al. (2012). “Detrimental effects of arachidonic acid and its metabolites in cellular and mouse models of Alzheimer’s disease: structural insight.” Neurobiology of aging 33(4): 831.e821–831.e831.

Andrade-Guerrero, J., et al. (2023). “Alzheimer’s Disease: An Updated Overview of Its Genetics.” International Journal of Molecular Sciences 24(4): 3754.

Ashley, E. A. (2016). “Towards precision medicine.” Nature Reviews Genetics 17(9): 507–522.

Basu, M., et al. (2021). “Predicting tissue-specific gene expression from whole blood transcriptome.” Science Advances 7(14): eabd6991.

Belfiori-Carrasco, L. F., et al. (2017). “A novel genetic screen identifies modifiers of age-dependent amyloid β toxicity in the Drosophila brain.” Frontiers in aging neuroscience 9: 61.

Berton, L., et al. (2015). Link prediction in graph construction for supervised and semi-supervised learning. 2015 International Joint Conference on Neural Networks (IJCNN), IEEE.

Camargo, A., et al. (2008). “Permutation–based statistical tests for multiple hypotheses.” Source code for biology and medicine 3: 1–8.

Chen, F., et al. (2021). “Dysfunction of the SNARE complex in neurological and psychiatric disorders.” Pharmacological Research 165: 105469.

Choi, S. H., et al. (2017). “Evaluation of logistic regression models and effect of covariates for case–control study in rna-seq analysis.” BMC bioinformatics 18(1): 1–13.

Consortium, E. (2015). “A roadmap for precision medicine in the epilepsies.” The Lancet Neurology 14(12): 1219–1228.

Cunnane, S. C., et al. (2012). “Plasma and brain fatty acid profiles in mild cognitive impairment and Alzheimer’s disease.” Journal of Alzheimer’s Disease 29(3): 691–697.

de Leeuw, F. A., et al. (2017). “Blood-based metabolic signatures in Alzheimer’s disease.” Alzheimer’s & Dementia: Diagnosis, Assessment & Disease Monitoring 8: 196–207.

Dos Santos, S. M., et al. (2019). “Mitochondrial dysfunction and alpha-lipoic acid: beneficial or harmful in Alzheimer’s disease?” Oxidative medicine and cellular longevity 2019.

Floudas, C. S., et al. (2014). “Identifying genetic interactions associated with late-onset Alzheimer’s disease.” BioData mining 7(1): 1–19.

Forssell, L. G., et al. (1989). “Early Stages of Late Onset Alzheimer’s Disease: II. Derangements in Protein Metabolism with Special Reference to Tryptophan, Tyrosine and Cystine.” Acta Neurologica Scandinavica 79(S121): 27–42.

Gamazon, E. R., et al. (2015). “A gene-based association method for mapping traits using reference transcriptome data.” Nature genetics 47(9): 1091–1098.

Golub, T. R., et al. (1999). “Molecular classification of cancer: class discovery and class prediction by gene expression monitoring.” Science 286(5439): 531–537.

Griffin, J. W. and P. C. Bradshaw (2017). “Amino acid catabolism in Alzheimer’s disease brain: friend or foe?” Oxidative medicine and cellular longevity 2017.

Günzel, D. and A. S. Yu (2013). “Claudins and the modulation of tight junction permeability.” Physiological reviews 93(2): 525–569.

Halloran, J. W., et al. (2015). “Prediction of the gene expression in normal lung tissue by the gene expression in blood.” BMC medical genomics 8(1): 1–6.

Hargis, K. E. and E. M. Blalock (2017). “Transcriptional signatures of brain aging and Alzheimer’s disease: What are our rodent models telling us?” Behavioural brain research 322: 311–328.

Hisano, R. (2018). Semi-supervised graph embedding approach to dynamic link prediction. Complex Networks IX: Proceedings of the 9th Conference on Complex Networks CompleNet 2018 9, Springer.

Hohman, T. J., et al. (2015). “The role of vascular endothelial growth factor in neurodegeneration and cognitive decline: exploring interactions with biomarkers of Alzheimer disease.” JAMA neurology 72(5): 520–529.

Hong, C., et al. (2020). “TRP channels as emerging therapeutic targets for neurodegenerative diseases.” Frontiers in Physiology 11: 238.

Hornik, K., et al. (1989). “Multilayer feedforward networks are universal approximators.” Neural networks 2(5): 359–366.

Hu, J., et al. (2010). “Computational analysis of tissue-specific gene networks: application to murine retinal functional studies.” Bioinformatics 26(18): 2289–2297.

Hu, Y. T., et al. (2019). “Early growth response-1 regulates acetylcholinesterase and its relation with the course of Alzheimer’s disease.” Brain Pathology 29(4): 502–512.

Jansen, I. E., et al. (2019). “Genome-wide meta-analysis identifies new loci and functional pathways influencing Alzheimer’s disease risk.” Nature genetics 51(3): 404–413.

Kalaria, R., et al. (1998). “Vascular endothelial growth factor in Alzheimer’s disease and experimental cerebral ischemia.” Molecular brain research 62(1): 101–105.

Kanehisa, M. (2002). The KEGG database. ‘In silico’simulation of biological processes: Novartis Foundation Symposium 247, Wiley Online Library.

Kaur, D., et al. (2021). “Decrypting the potential role of α-lipoic acid in Alzheimer’s disease.” Life Sciences 284: 119899.

Langfelder, P. and S. Horvath (2008). “WGCNA: an R package for weighted correlation network analysis.” BMC bioinformatics 9(1): 1–13.

Lee, A. Y., et al. (2018). “Alpha-linolenic acid regulates amyloid precursor protein processing by mitogen-activated protein kinase pathway and neuronal apoptosis in amyloid beta-induced SH-SY5Y neuronal cells.” Applied Biological Chemistry 61: 61–71.

Li, Y., et al. (2021). “Genomics of Alzheimer’s disease implicates the innate and adaptive immune systems.” Cellular and Molecular Life Sciences: 1–30.

Liu, P., et al. (2021). “Phenylalanine metabolism is dysregulated in human hippocampus with Alzheimer’s disease related pathological changes.” Journal of Alzheimer’s Disease 83(2): 609–622.

Lonsdale, J., et al. (2013). “The genotype-tissue expression (GTEx) project.” Nature genetics 45(6): 580–585.

Lu, L., et al. (2021). “Learning nonlinear operators via DeepONet based on the universal approximation theorem of operators.” Nature machine intelligence 3(3): 218–229.

Lu, R., et al. (2017). “TRPC channels and Alzheimer’s disease.” Transient Receptor Potential Canonical Channels and Brain Diseases: 73–83.

Maglott, D., et al. (2010). “Entrez Gene: gene-centered information at NCBI.” Nucleic acids research 39(suppl_1): D52–D57.

Maniatis, T., et al. (1987). “Regulation of inducible and tissue-specific gene expression.” Science 236(4806): 1237–1245.

Margiotta, A. (2021). “Role of SNAREs in neurodegenerative diseases.” Cells 10(5): 991.

Martínez, M. and N. C. Inestrosa (2021). “The transcriptional landscape of Alzheimer’s disease and its association with Wnt signaling pathway.” Neuroscience & Biobehavioral Reviews 128: 454–466.

Maruszak, A. and S. Thuret (2014). “Why looking at the whole hippocampus is not enough—a critical role for anteroposterior axis, subfield and activation analyses to enhance predictive value of hippocampal changes for Alzheimer’s disease diagnosis.” Frontiers in cellular neuroscience 8: 95.

McGeer, P., et al. (1992). “Distribution of clusterin in Alzheimer brain tissue.” Brain research 579(2): 337–341.

Mercola, J. and C. R. D’Adamo (2023). “Linoleic acid: A narrative review of the effects of increased intake in the standard American diet and associations with chronic disease.” Nutrients 15(14): 3129.

Mez, J., et al. (2017). “Two novel loci, COBL and SLC10A2, for Alzheimer’s disease in African Americans.” Alzheimer’s & dementia 13(2): 119–129.

Mirza, A., et al. (2017). “Aluminium in brain tissue in familial Alzheimer’s disease.” Journal of Trace Elements in Medicine and Biology 40: 30–36.

Montine, T. J. and J. D. Morrow (2005). “Fatty acid oxidation in the pathogenesis of Alzheimer’s disease.” The American journal of pathology 166(5): 1283–1289.

Morash, M., et al. (2018). “The role of next-generation sequencing in precision medicine: a review of outcomes in oncology.” Journal of personalized medicine 8(3): 30.

Mueller, S. G., et al. (2005). “The Alzheimer’s disease neuroimaging initiative.” Neuroimaging Clinics 15(4): 869–877.

Ong, C.-T. and V. G. Corces (2011). “Enhancer function: new insights into the regulation of tissue-specific gene expression.” Nature Reviews Genetics 12(4): 283–293.

Patel, D., et al. (2019). “Association of rare coding mutations with Alzheimer disease and other dementias among adults of European ancestry.” JAMA network open 2(3): e191350–e191350.

Pereira, C. D., et al. (2018). “ABC transporters are key players in Alzheimer’s disease.” Journal of Alzheimer’s Disease 61(2): 463–485.

Pérez-González, M., et al. (2020). “PLA2G4E, a candidate gene for resilience in Alzheimer s disease and a new target for dementia treatment.” Progress in neurobiology 191: 101818.

Romanitan, M. O., et al. (2010). “Altered expression of claudin family proteins in Alzheimer’s disease and vascular dementia brains.” Journal of cellular and molecular medicine 14(5): 1088–1100.

Sabaie, H., et al. (2023). “Expression analysis of inhibitory B7 family members in Alzheimer’s disease.” Metabolic Brain Disease 38(8): 2563–2572.

Saura, C. A. and J. Valero (2011). “The role of CREB signaling in Alzheimer’s disease and other cognitive disorders.”

Saxena, S., et al. (2021). “Structural and functional analysis of disease-associated mutations in GOT1 gene: An in silico study.” Computers in Biology and Medicine 136: 104695.

Schedin-Weiss, S., et al. (2017). “Monoamine oxidase B is elevated in Alzheimer disease neurons, is associated with γ-secretase and regulates neuronal amyloid β-peptide levels.” Alzheimer’s Research & Therapy 9: 1–19.

Schwab, C., et al. (2000). “Casein kinase 1 delta is associated with pathological accumulation of tau in several neurodegenerative diseases.” Neurobiology of aging 21(4): 503–510.

Snowden, S. G., et al. (2017). “Association between fatty acid metabolism in the brain and Alzheimer disease neuropathology and cognitive performance: A nontargeted metabolomic study.” PLoS medicine 14(3): e1002266.

Spulber, S., et al. (2012). “Claudin expression profile separates Alzheimer’s disease cases from normal aging and from vascular dementia cases.” Journal of the neurological sciences 322(1-2): 184–186.

Su, J., et al. (1992). “Localization of heparan sulfate glycosaminoglycan and proteoglycan core protein in aged brain and Alzheimer’s disease.” Neuroscience 51(4): 801–813.

Subramanian, A., et al. (2005). “Gene set enrichment analysis: a knowledge-based approach for interpreting genome-wide expression profiles.” Proceedings of the National Academy of Sciences 102(43): 15545–15550.

Tanaka, S., et al. (1989). “Tissue-specific expression of three types of β-protein precursor mRNA: enhancement of protease inhibitor-harboring types in Alzheimer’s disease brain.” Biochemical and biophysical research communications 165(3): 1406–1414.

Thomas, M. H. and J. L. Olivier (2016). “Arachidonic acid in Alzheimer’s disease.” Journal of Neurology & Neuromedicine 1(9).

Uzor, N.-E., et al. (2020). “Peroxisomal dysfunction in neurological diseases and brain aging.” Frontiers in cellular neuroscience 14: 44.

Van Horssen, J., et al. (2003). “Heparan sulphate proteoglycans in Alzheimer’s disease and amyloid-related disorders.” The Lancet Neurology 2(8): 482–492.

Vaswani, A., et al. (2017). “Attention is all you need.” Advances in neural information processing systems 30.

Veličković, P., et al. (2017). “Graph attention networks.” arXiv preprint arXiv:1710.10903.

Vilgis, S. and H.-P. Deigner (2018). Sequencing in precision medicine. Precision medicine, Elsevier: 79–101.

Wang, T., et al. (2021). “A pipeline for RNA-seq based eQTL analysis with automated quality control procedures.” BMC bioinformatics 22(9): 1–18.

Watanabe, T., et al. (2012). “Neuronal expression of F-box and leucine-rich-repeat protein 2 decreases over Braak stages in the brains of Alzheimer’s disease patients.” Neurodegenerative Diseases 11(1): 1–12.

Whitehead, A. and D. L. Crawford (2005). “Variation in tissue-specific gene expression among natural populations.” Genome biology 6(2): 1–14.

Wissmann, P., et al. (2013). “Immune activation in patients with Alzheimer’s disease is associated with high serum phenylalanine concentrations.” Journal of the neurological sciences 329(1-2): 29–33.

Wu, L., et al. (2022). Graph neural networks: foundation, frontiers and applications. Proceedings of the 28th ACM SIGKDD Conference on Knowledge Discovery and Data Mining.

Xu, W., et al. (2020). “Blood-based multi-tissue gene expression inference with Bayesian ridge regression.” Bioinformatics 36(12): 3788–3794.

Yasojima, K., et al. (2000). “Casein kinase 1 delta mRNA is upregulated in Alzheimer disease brain.” Brain research 865(1): 116–120.

Zhao, Y., et al. (2021). “Targeted metabolomics study of early pathological features in hippocampus of triple transgenic Alzheimer’s disease male mice.” Journal of Neuroscience Research 99(3): 927–946.

Zhou, J., et al. (2020). “Graph neural networks: A review of methods and applications.” AI open 1: 57–81.

Zhu, Q.-B., et al. (2016). “MicroRNA-132 and early growth response-1 in nucleus basalis of Meynert during the course of Alzheimer’s disease.” Brain 139(3): 908–921.

