## Supplemental Materials for "Cross-tissue Graph Attention Networks for Semi-supervised Gene Expression Prediction"

A. **Implementation details**

In this section we will introduce details of implementation of our method. We use packages in R 4.3.3 to construct gene co-expression network for each tissue. Then we leverage packages with Python 3.8.8 for modeling.

***Gene co-expression network construction***

We employ the package “WGCNA” (v1.72.5) in R with default parameter settings for gene co-expression network construction.

***Source encoding***

Source encoding process uses four-layer GATs by “GATConv” of DGL 2.2.1 in Python, to encode the network of the source tissue. The detailed parameters in GATs are listed below:

| # layer | #in_channel | #out_channel | #head |
| --- | --- | --- | --- |
| 1 | 1 | 1024 | 8 |
| 2 | 1024 x 8 | 1024 | 1 |
| 3 | ELU | | |
| 4 | 1024 | 1024 | 8 |
| 5 | 1024 x 8 | 1024 | 1 |
| 6 | ELU | | |
| 7 | 1024 | 1024 | 8 |
| 8 | 1024 x 8 | 1024 | 1 |
| 9 | ELU | | |
| 10 | 1024 | 1024 | 8 |
| 11 | 1024 x 8 | 1024 | 1 |
| 12 | ELU | | |

Details of MLP layers are as below:

| # layer | #in | #out |
| --- | --- | --- |
| 1 | 1028 | 512 |
| 2 | ELU | |
| 3 | 512 | 256 |
| 4 | ELU | |
| 5 | 256 | 256 |
| 6 | ELU | |
| 7 | 256 | 256 |
| 8 | ELU | |
| 9 | 256 | 64 |
| 10 | ELU | |
| 11 | 64 | 16 |
| 12 | ELU | |
| 13 | 16 | 4 |
| 14 | ELU | |
| 15 | 4 | 1 |

***Target decoding***

Link prediction layers involve two-layer GAT followed by three-layer MLP. The details of GAT layers are as follows:

| # layer | #in_channel | #out_channel | #head |
| --- | --- | --- | --- |
| 1 | 1 | 1024 | 8 |
| 2 | 1024 x 8 | 1024 | 1 |
| 3 | ELU | | |
| 4 | 1024 | 1024 | 8 |
| 5 | 1024 x 8 | 1024 | 1 |
| 6 | ELU | | |

MLP layers are as follows:

| # layer | #in | #out |
| --- | --- | --- |
| 1 | 1028 | 128 |
| 2 | ELU | |
| 3 | 128 | 128 |
| 4 | ELU | |
| 5 | 128 | 1 |

Similarlly, the predictor also involves two-GAT and eight-layer MLP. Details of GAT layers are as follows:

| # layer | #in_channel | #out_channel | #head |
| --- | --- | --- | --- |
| 1 | 1 | 1024 | 8 |
| 2 | 1024 x 8 | 1024 | 1 |
| 3 | ELU | | |
| 4 | 1024 | 1024 | 8 |
| 5 | 1024 x 8 | 1024 | 1 |
| 6 | ELU | | |

MLP layers are as follows:

| # layer | #in | #out |
| --- | --- | --- |
| 1 | 1028 | 512 |
| 2 | ELU | |
| 3 | 512 | 256 |
| 4 | ELU | |
| 5 | 256 | 256 |
| 6 | ELU | |
| 7 | 256 | 256 |
| 8 | ELU | |
| 9 | 256 | 64 |
| 10 | ELU | |
| 11 | 64 | 16 |
| 12 | ELU | |
| 13 | 16 | 4 |
| 14 | ELU | |
| 15 | 4 | 1 |

B. **Compare best-predicted genes among tissues**

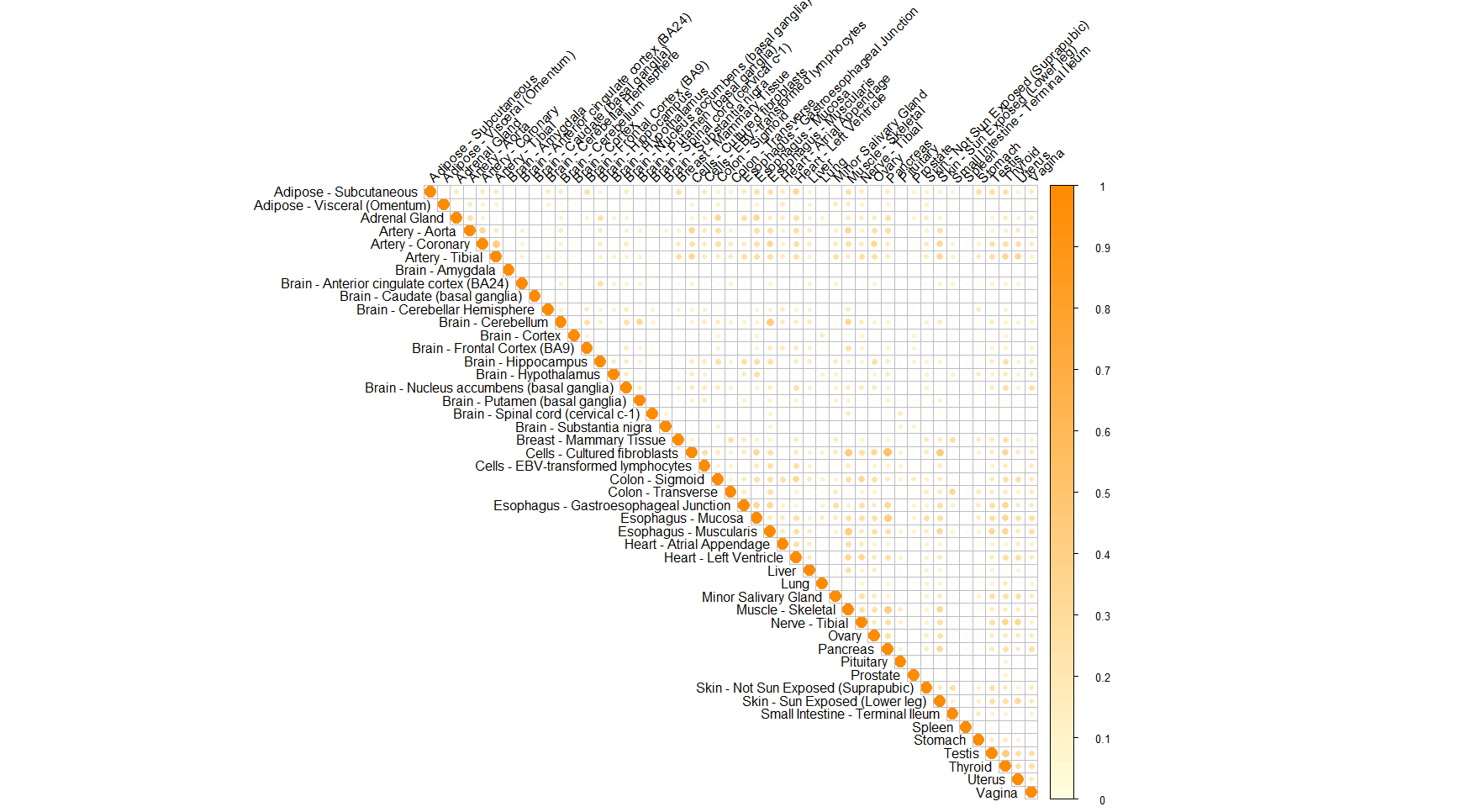

**Figure S1:** The overlap among the top ten predicted genes (based on PCC) in all tissue pairs in terms of Jaccard index.

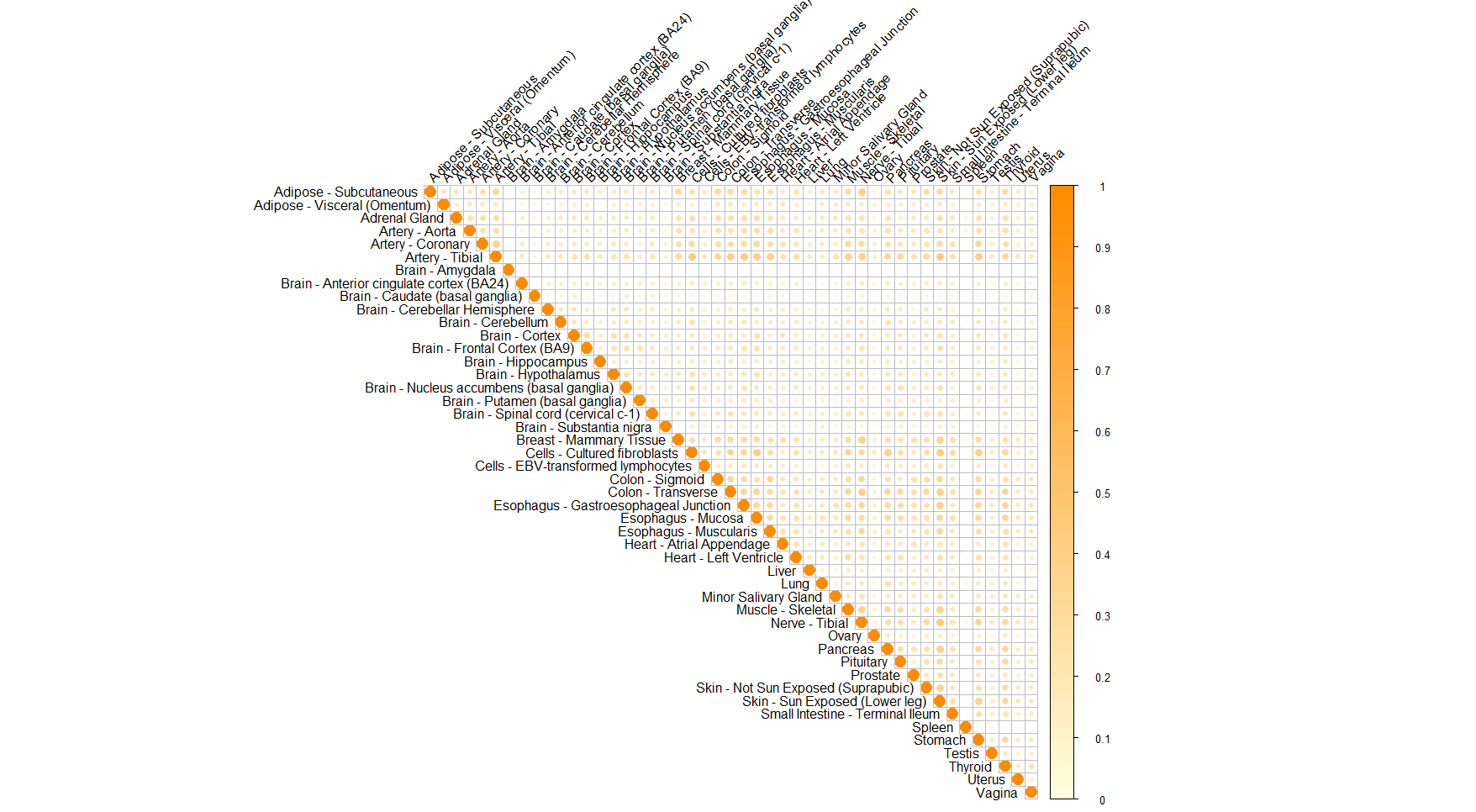

**Figure S2:** The overlap among the top 100 predicted genes (based on PCC) in all tissue pairs in terms of Jaccard index.

C. **Prediction quality of all genes in each tissue.**

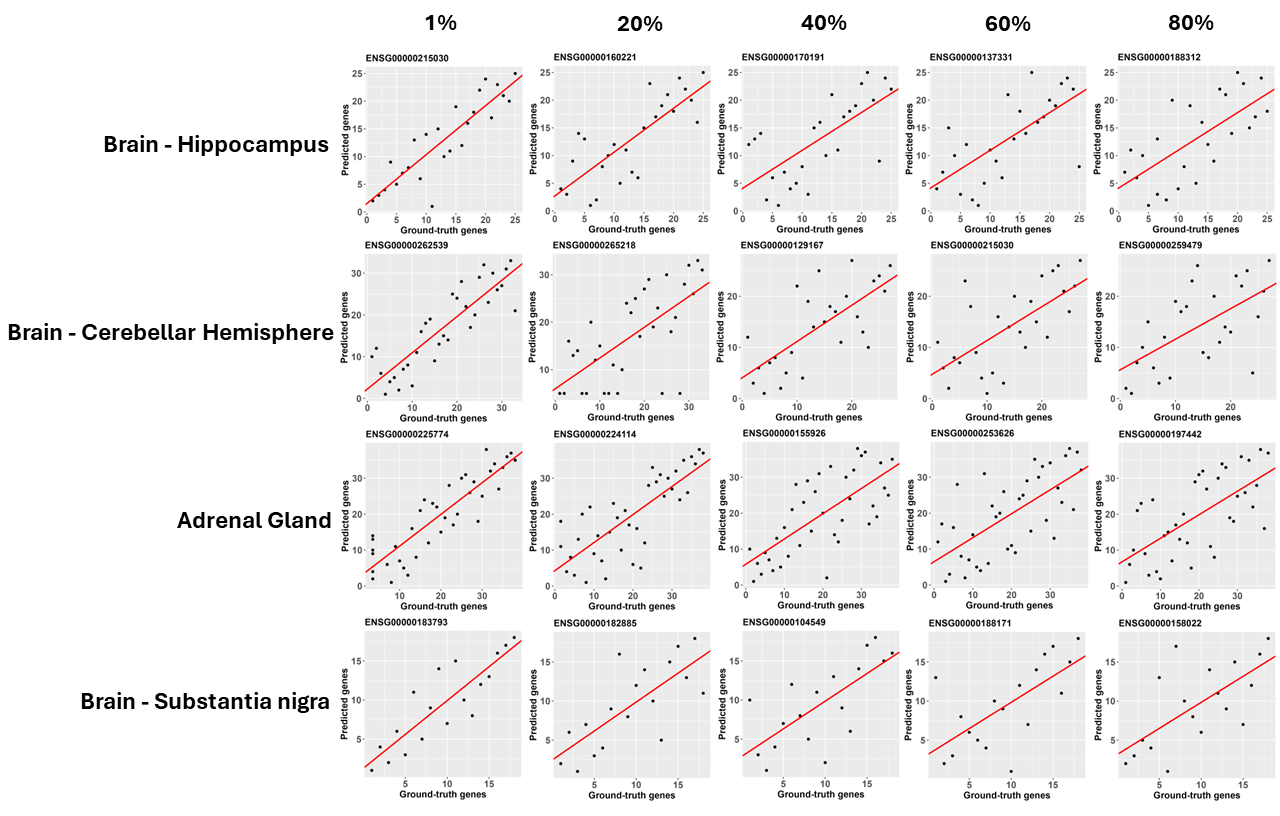

**Figure S3:** Compare predicted values against true expressions for genes at 1%, 20%, 40%, 60% and 80% ranked by PCC in four best predicted tissues (i.e., Brain Hippocampus, Brain Cerebellar Hemisphere, Adrenal Gland and Brain Substantia Nigra). Red line is the linear regression line fitted used predicted and true values.

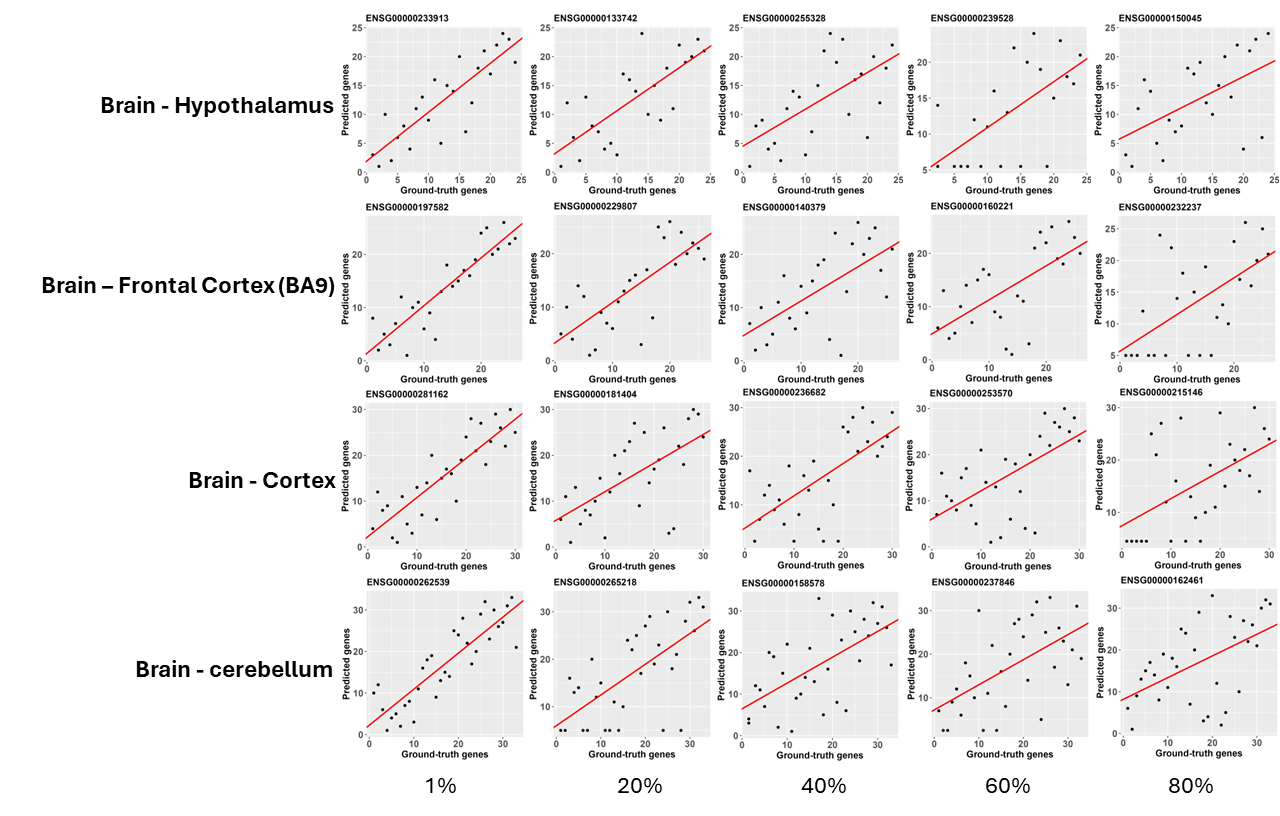

**Figure S4:** Compare predicted values against true expressions for genes at 1%, 20%, 40%, 60% and 80% ranked by PCC in four worst predicted tissues (i.e., Brain Hypothalamus, Brain Frontal Cortex (BA9), Brain Cortex and Brain Cerebellum). Red line is the linear regression line fitted used predicted and true values.

For each gene, we rank their prediction performance in each tissue based on PCC. As we have 47 tissues in total, the ranking of each gene ranges from 1 to 47. The results are saved in the file “gene_rank_by_tissue.csv”.

D. **Application to identify disease-associated genes**

In this session, we provide more evidence to verify AD-associated genes predicted by C2-GAT. Based on Table 1, *HHLA2* (p-value=$1.01\times{10}^{-3}$) belongs to B7 family, whose members were found to have significantly higher in AD patients. The expression level of *IGSF6* (p-value=$1.27\times{10}^{-3}$) was found to be proportional to AD progresses(Hargis and Blalock 2017). *SCIMP* (p-value$=1.32\times{10}^{-3}$) is identified to be associated with alteration in TLR-mediated microglial phagocytosis via MHC II(Andrade-Guerrero, Santiago-Balmaseda et al. 2023). AD dementia correlates with increased MHC II+ microglia-mediated immunity(Li, Laws et al. 2021). Kalaria, Cohen et al. (1998) find that enhanced *VEGF* (p-value$=1.36\times{10}^{-3}$) immunoreactivity in clusters of reactive astrocytes in the neocortex of subjects with AD compared to elderly controls. The last-ranked gene, *WLS* (p-value$=1.67\times{10}^{-3}$), also called Wnt ligand secretion mediation, is essential for both the developing and adult central nervous systems(Aghaizu, Jin et al. 2020). AD exhibits a range of pathophysiological manifestations including aberrant amyloid precursor protein processing, tau pathology, synapse loss, neuroinflammation and blood brain barrier breakdown, which have been associated to a greater or lesser degree with abnormal *Wnt* signaling(Aghaizu, Jin et al. 2020). Therefore, all top AD related genes identified by the differential expression analysis are supported by the existing scientific findings, indicating the effectiveness of gemGAT in preserving the distribution of predicted genes in different populations (e.g., AD patients and healthy population).

**Table S1:** Top ten AD-associated genes in Whole Blood based on logistic-regression p-values.

| **Gene symbol** | **p-value** | **Permutation p-value** |
| --- | --- | --- |
| *ASB2* | $2.71\times{10}^{-4}$ | $4.00\times{10}^{-4}$ |
| *CREB5* | $4.64\times{10}^{-4}$ | $6.00\times{10}^{-4}$ |
| *DNAAF11* | $5.85\times{10}^{-4}$ | $4.00\times{10}^{-4}$ |
| *FAMI85A* | $7.59\times{10}^{-4}$ | $4.00\times{10}^{-3}$ |
| *FBXL13* | $8.39\times{10}^{-4}$ | $4.00\times{10}^{-4}$ |
| *GTSE1* | $9.31\times{10}^{-4}$ | $1.00\times{10}^{-3}$ |
| *IGSF6* | $1.09\times{10}^{-3}$ | $4.00\times{10}^{-4}$ |
| *TENM1* | $1.17\times{10}^{-3}$ | $8.00\times{10}^{-4}$ |
| *TRPC3* | $1.30\times{10}^{-3}$ | $1.80\times{10}^{-3}$ |
| *WLS* | $1.34\times{10}^{-3}$ | $1.00\times{10}^{-3}$ |

As shown in Supplemental Table S1, For other genes, *CREB* (p-value$=4.64\times{10}^{-4}$) signaling is essential for long-lasting changes in synaptic plasticity that mediates the conversion of short-term memory to long-term memory(Saura and Valero 2011). The $\beta$-amyloid peptide, which plays a crucial role in the pathogenesis of AD, alters hippocampal-dependent synaptic plasticity and memory and mediates synapse loss through the CREB signaling pathway(Saura and Valero 2011). Additionally, Besides, *GTSE1* (p-value$=9.31\times{10}^{-4}$) and *TRPC3* (p-value$=1.30\times{10}^{-3}$) are also playing significant roles in the mechanism of AD(Lu, He et al. 2017, Patel, Mez et al. 2019, Hong, Jeong et al. 2020)

E. **Application to disease-associated pathways**

As shown in **Table 2,** Arachidonic acid metabolism (p-value$=1.71\times{10}^{-2}$) are found to be involved in Alzheimer’s disease through several mechanisms, such as neuroinflammation and synaptic functions(Thomas and Olivier 2016). Differences in plasma fatty acid (p-value$=2.96\times{10}^{-2}$) profiles between AD, mild cognitive impairment, and those with no cognitive impairment are found apparent(Cunnane, Schneider et al. 2012). ABC transporters (p-value$=3.14\times{10}^{-2}$), localized on the surface of brain endothelial cells of the blood brain barrier and brain parenchyma, have altered function in the pathogenesis and progression of AD(Abuznait and Kaddoumi 2012). Inhibition of the formation of the SNARE complex, defects in the SNARE-dependent exocytosis and altered regulation of SNARE-mediated vesicle fusion have been associated with neurodegeneration(Margiotta 2021).

The top-ranked pathways in the whole blood are shown in **Table S2**.

**Table S2:** Top ten AD-associated pathways identified in Whole Blood based on logistic-regression p-values.

| **Pathway** | **NES** | **p-value** |
| --- | --- | --- |
| retinol metabolism | 1.58 | $4.11\times{10}^{-3}$ |
| dorso ventral axis formation | 1.40 | $5.57\times{10}^{-2}$ |
| proximal tubule bicarbonate reclamation | 1.37 | $6.59\times{10}^{-2}$ |
| citrate cycle tca cycle | 1.37 | $5.23\times{10}^{-2}$ |
| arginine and proline metabolism | 1.34 | $3.79\times{10}^{-2}$ |
| proteasome | 1.33 | $5.26\times{10}^{-2}$ |
| fatty acid metabolism | 1.30 | $7.95\times{10}^{-2}$ |
| endometrial cancer | 1.26 | $8.89\times{10}^{-2}$ |
| fructose and mannose metabolism | 1.23 | $1.43\times{10}^{-1}$ |
| notch signaling pathway | 1.22 | $1.32\times{10}^{-1}$ |

F. **Comparison with regression-based methods**

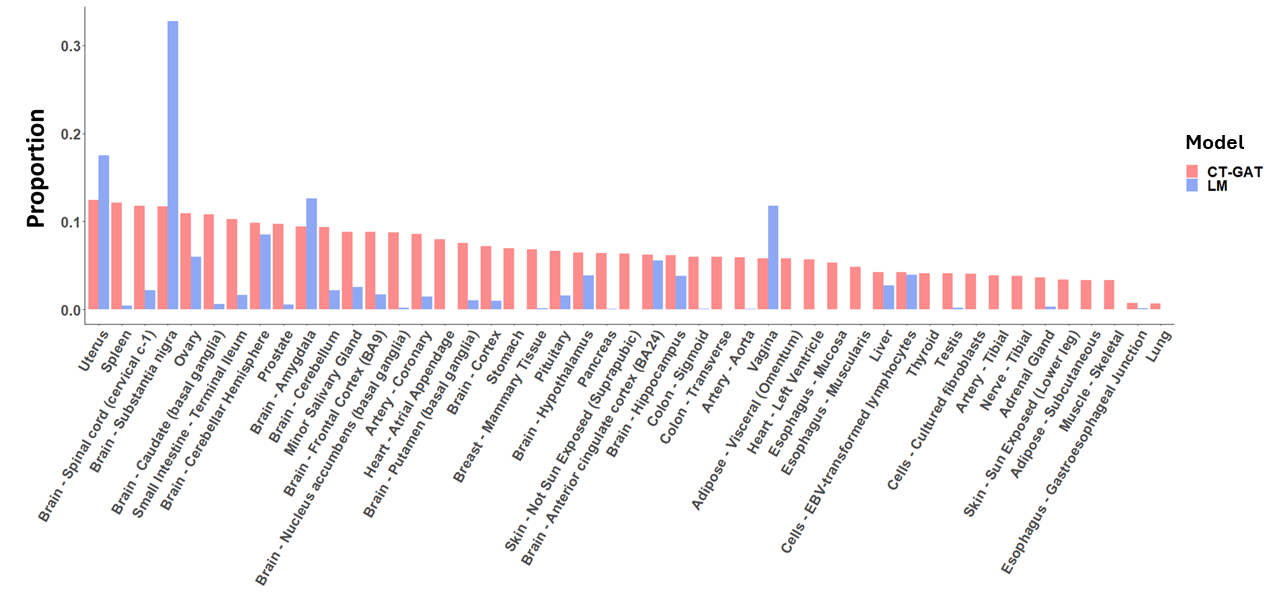

**Figure S5:** Proportion of well-predicted genes in terms of distributional similarities (i.e., with p-value > 0.05 in Kolmogorov–Smirnov test) between their predicted and true expressions using gemGAT and TEEBOT. Note that PrediXcan is not shown in B and C as no well-predicted genes are identified.

G. **Graph attention networks.**

We explicitly present the formulation of the layer of GATs(Veličković, Cucurull et al. 2017) in this section. Suppose $\mathcal{G}$ is the graph representing interactions among nodes $\left\{ v_{i}\mathcal{\in V} \right\}_{i=1}^{N}$ (e.g., genes in our case), $\mathcal{G}$ can also be written into an adjacency matrix $A$, which as the following form:

$$A_{ij}=\left\{ \begin{aligned} 1, v_{i} interacts with v_{j} \\ 0, {no interaction between v}_{i} and v_{j} \end{aligned} \right.$$

Suppose $\mathcal{X=}\left\{ x_{i} \right\}_{i=1}^{N}$ is the set of features of $N$ nodes in $\mathcal{G}$, with $x_{i}\in\mathbb{R}^{d}$. To compute $GAT(\mathcal{G,X;}\beta)$, where $\beta$ is the set of model parameters, we concatenate all feature vectors, $x_{i}, i=1, 2, \ldots, N$, in $\mathcal{X}$ to form $X\in\mathbb{R}^{N\times d}$, which is the feature matrix of $N$ nodes in $\mathcal{G}$. We learn the attention mechanism (i.e., $\alpha_{ij}$ for the edge that connects $v_{i}$ and $v_{j}$) via a single-layer feedforward neural network parameterized by a weight vector $a\in\mathbb{R}^{2d}$:

$$\alpha_{ij}=\frac{\exp(LeakyReLU(a^{T}[Wx_{i}\|Wx_{j}]))}{\sum_{k\in\mathcal{N}_{i}} \exp(LeakyReLU(a^{T}[Wx_{i}\|Wx_{k}]))},$$

where $\cdot^{T}$ represents transposition, $\|$ is the concatenation operation and both $W$, $a$ are learnable parameters. $LeakyReLU(\cdot)$ is the activation function to add nonlinearity to the layer. Once obtained, these normalized attention coefficients are used to compute a linear combination of the features corresponding to them, to serve as the final output features for every node, after applying a nonlinearity:

$$x_{i}^{'}=\sigma\left( \sum_{j\in\mathcal{N}_{i}} \alpha_{ij}Wx_{j} \right),$$

where $\sigma$ is an activation function to add nonlinearity such as LeakyReLU, ReLU, Tanh, etc. In practice, we borrow a $K$ multi-head attention following Vaswani, Shazeer et al. (2017):

$$x_{i}^{'}=\left. \right\|_{k=1}^{K}\sigma\left( \sum_{j\in\mathcal{N}_{i}} \alpha_{ij}^{k}W^{k}x_{j} \right),$$

where $\alpha_{ij}^{k}$ and $W^{k}$ are the learnable attention coefficient and linear coefficient respectively for the head $k$. $x_{i}'$ is the output of GATs for node $v_{i}\mathcal{\in V}$, and model parameters $\beta=\{a^{k}, W^{k}: k=1,2,\ldots,K\}$.

H. **Multilayer perceptron.**

In this section we present the formulation of the MLP layer, $MLP(\mathcal{X,}\beta)$, where $\mathcal{X}$ is the set of features $\mathcal{X=}\left\{ x_{i} \right\}_{i=1}^{N}$ with $x_{i}\in\mathbb{R}^{d}$. We further suppose $X\in\mathbb{R}^{N\times d}$ as the concatenation of all features in $\mathcal{X}$. With the input $X$, the output of MLP is $X^{'}$ formulated as follows:

$$X^{'}=\sigma\left( WX \right),$$

where $W$ is the learnable parameter, and $\sigma(\cdot)$ is the activation function to add nonlinearity to the layer, such as ReLU, ELU, Tanh, etc.
